# Convergent genomic signatures of adaptation to an extreme environment in cyprinoid fishes

**DOI:** 10.1101/2022.03.30.486405

**Authors:** Chao Tong, Miao Li

## Abstract

Cyprinoids are a worldwide distributed and diverse group of freshwater fish with more than 3,000 species. Although primarily freshwater, some cyprinoid species had convergently evolved to thrive in extreme environments and underlying genetic mechanisms remain unclear. Here, we leveraged 32 cyprinoid fish genomes to identify common genomic changes associated with convergent adaptation to highly saline and alkaline water in two East Asian cyprinoid fish species, *Gymnocypris przwalskii* and *Leuciscus waleckii*, representing two independent extremophile fish lineages. We found that genome-wide rate of nonsynonymous substitution and signal of intensified selection is higher in extremophile relative to non-extremophile fish taxa. We further tested gene-wide molecular convergence and found hundreds of genes tended to experience convergent shifts in selections in extremophile fish taxa, including convergent acceleration and positive selection. These genes were associated with several key functions, such as nervous system development, reproduction, ion transport and immune response, and included genes that previously have been implicated for saline or alkaline tolerance in fish. Additionally, comparative transcriptomic analyses defined the convergent roles of differentially expressed genes under selection in extremophile fish taxa during convergent adaptation. Taken together, our work provides insights into the genomic basis of convergent adaptation to extreme environments in fish.

## Introduction

Convergent evolution is especially common in organisms that have independently adapted to the same environments (Stern 2013; Storz 2016). That is the repeated evolution of similar phenotypic innovations in widely divergent evolutionary lineages. A major motivation in evolutionary biology is to determine the genetic changes that underlie phenotypic convergent evolution. In several known cases, selection repeatedly targets the same genes, such as Prestin in echolocation in bats and marine mammals (Li *et al*. 2010; Liu *et al*. 2010), Nav1.4 in toxin resistance in snakes (Feldman *et al*. 2012), and AgRP2 in stripe patterns in cichlid fishes (Kratochwil *et al*. 2018). A key feature of these phenotypic convergences is that identical amino acid (AA) substitutions within the same protein-coding gene occurred independently in unrelated taxa. In comparison to convergence in a single gene, numerous comparative genomics studies have identified massive convergent AA substitutions, that are widespread throughout the genome. Examples include, but are not limited to, echolocating mammals (Parker *et al*. 2013), marine mammals (Foote *et al*. 2015), flight-degenerate birds (Pan *et al*. 2019). It is increasingly clear that convergent evolution can also cause convergent shifts in evolutionary selective pressures, including nucleotide substitution rate (Yu *et al*.2016; Sun *et al*. 2018; Wu *et al*. 2020) and protein evolutionary rate (Partha *et al*. 2017; Kowalczyk *et al*. 2020). In addition, recent genome-wide studies suggest molecular convergence can result from concerted gene expression alteration in diverse groups of organisms adaptation to the same environmental stresses (Hao *et al*. 2019; Greenway *et al*. 2020). However, the degree to which convergent phenotypic outcomes result from the same genetic changes remains an unanswered question. Thus, new studies employing genome-wide approaches may provide insight into the signatures of convergent evolution in an explosion of available sequenced organisms (Sackton & Clark 2019).

Extremophiles form a fascinating system to study the convergent phenotypic evolution (Brown *et al*. 2019; Greenway *et al*. 2020; Xu *et al*. 2020; Yang *et al*. 2021). For example, the recent burst of fish genome data has enabled the expansion of molecular convergent analyses in several extremophile fish lineages, such as livebearing fishes convergently adapted to hydrogen sulfide (Brown *et al*. 2019; Greenway *et al*. 2020), highland fishes convergently adapted to hypoxia (Yang *et al*. 2021), while the underlying genomic mechanisms remain largely unexplored. Cyprinoids are a diverse family that included over 3,000 species with worldwide distributions (Winfield & Nelson 2012). Although cyprinoids occur primarily in freshwater habitats, some species can even thrive in extreme environment, such as *Gymnocypris przwalskii* and *Leuciscus waleckii* in hypersaline and hyperalkaline water, representing a striking example of convergent phenotypic evolution that has long interested biologists (Xu *et al*. 2017; Tong *et al*. 2017, 2021; Tong & Li 2020). Specifically, *G. przwalskii* (family: Cyprinidae) inhabits Lake Qinghai with high salinity (up to 13‰) and high alkalinity (up to pH 9.4) on the northeastern Tibetan Plateau, *L. waleckii* (family: Leuciscidae) survives in Lake Nali Nur with relatively high salinity (up to 6‰) and extreme alkalinity (up to pH 9.6) on the eastern Inner Mongolian Plateau (Fig. 1). Notably, both extremophile fish survive in freshwater of connective rivers during the spawning migration (Xu *et al*. 2017; Tong *et al*. 2017). For this, several recent genome-wide studies, including ours, have reported the evolutionary changes associated with hypersaline or hyperalkaline adaptations in a single extremophile fish relative to other phylogenetically distant teleost fish species (Xu *et al*. 2017; Tong *et al*. 2017, 2021; Tong & Li 2020). However, how phylogenetic background contributes to convergent adaptation to extremely saline or alkaline environments is largely unclear, in particular in a group of cyprinoid fishes.

**Figure 1.**
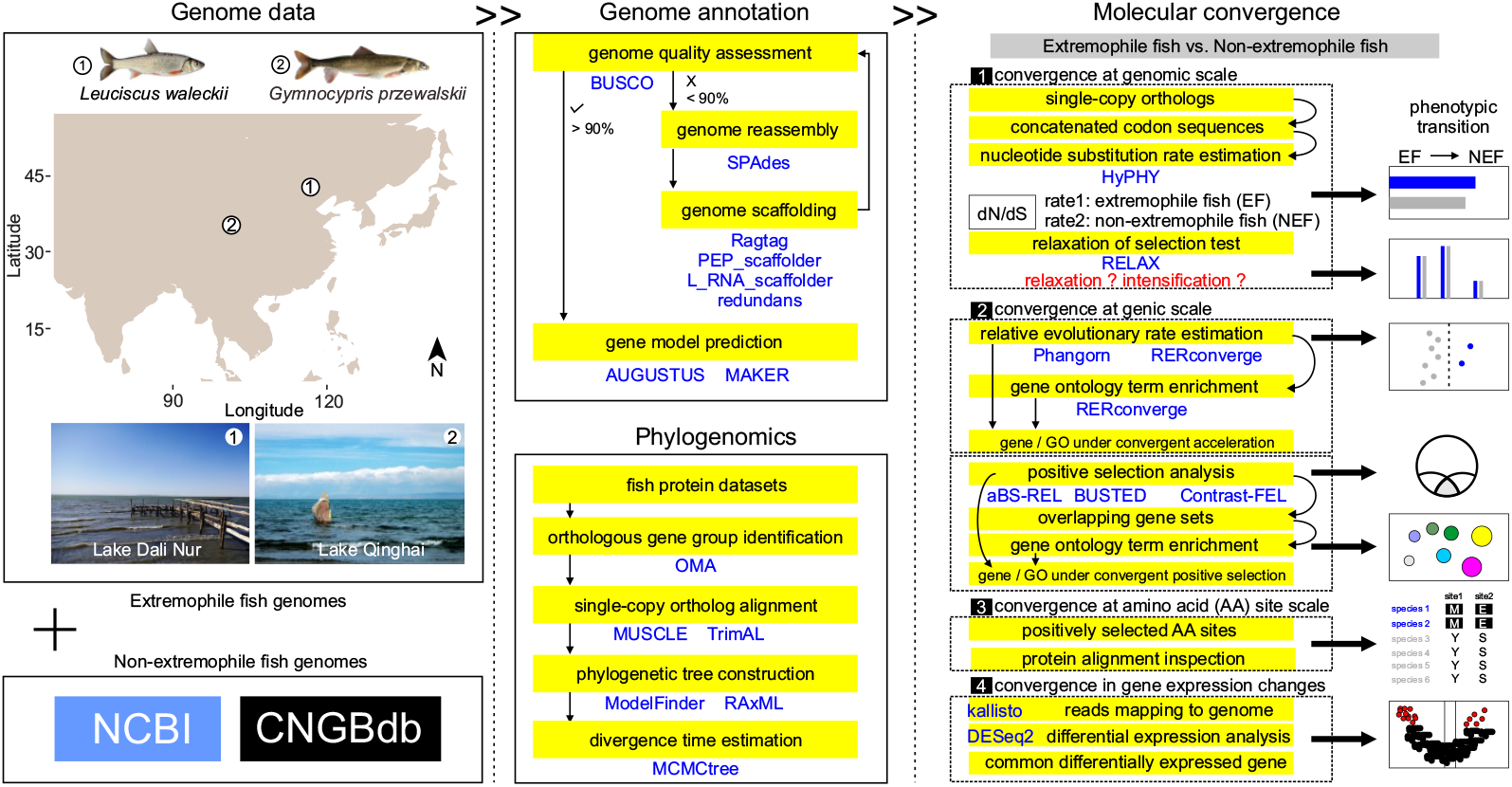
Sampling location, study design and workflow. Map showing the geographic locations of extremophile fish, *Leuciscus waleckii* from Lake Dali Nur and *Gymnocypris przewalskii* from Lake Qinghai in eastern Asia. Pictures of the extremophile fishes and their dwelling lakes (from top to bottom) (photo credit: Chao Tong, Xiaoxiao He). The schematic represents the analysis pipeline as follows: genome data collection, genome annotation, phylogenomics, and molecular convergence analyses. Technical details of the workflow are provided in the Materials and Methods section.

**Figure 2.**
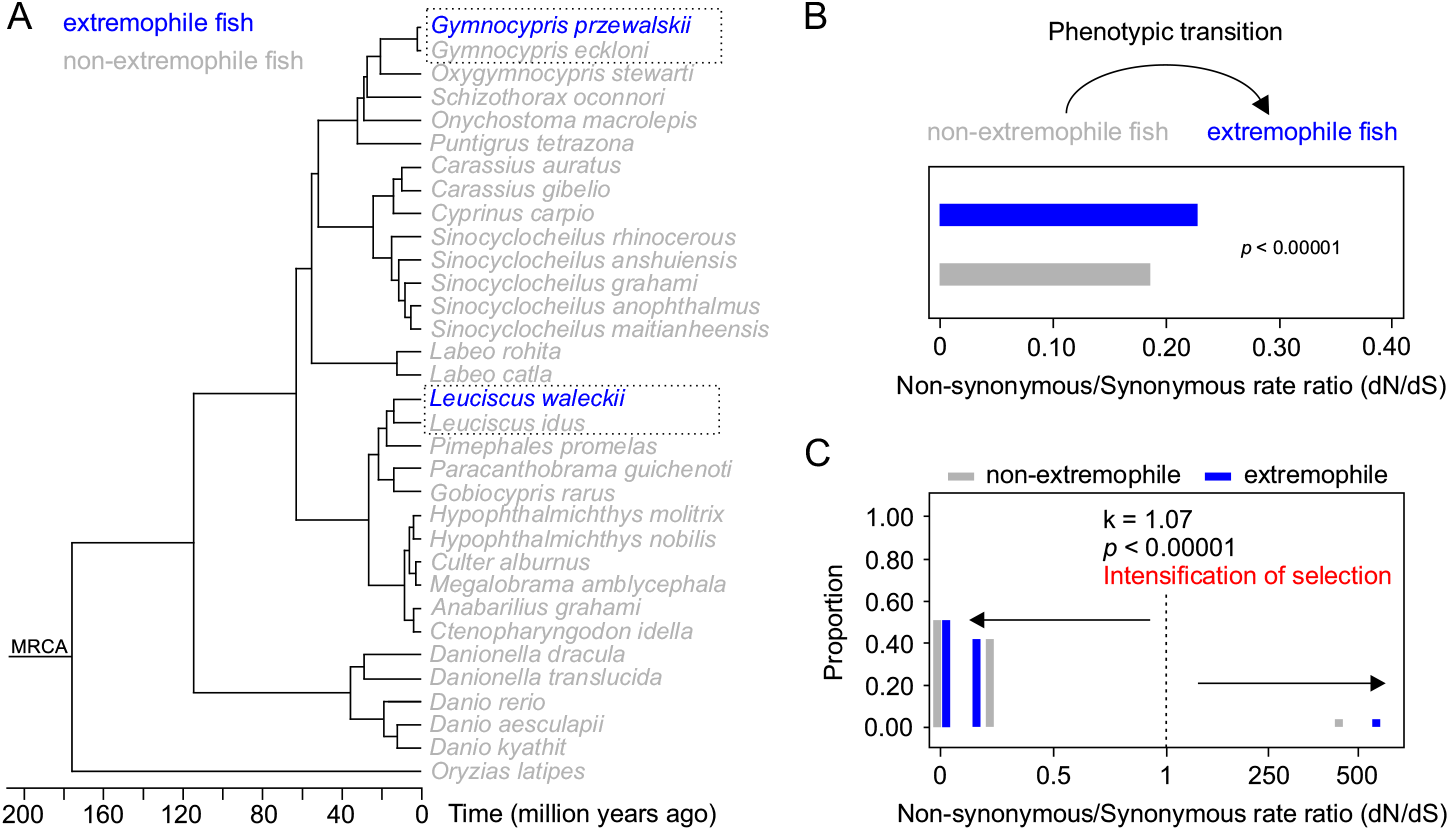
Genome-scale phylogeny of study cyprinoid fishes and genome-wide pattern of molecular evolution in extremophile fish taxa. (A) A maximum likelihood phylogenetic tree with estimated divergence time of the 32 cyprinoid fish species included in the study. Phylogenetic tree was inferred from 3,784 core shared single-copy orthologs, that was supported for all nodes with 100% bootstrap values. (B) Bar plot depicting the genome-wide rates of molecular evolution (also known as dN/dS) during the convergent phenotypic transition from non-extremophile to extremophile in fish. (C) Multi-bar plot depicting the pattern of intensified selection (k > 1) experienced across sites in genomes of extremophile relative to non-extremophile fish taxa.

Our study aimed to determine whether there are statistically supported common genomic signatures associated with convergent adaptation to extreme environment in cyprinoid fish. We leveraged a newly reassembled and 31 relevant available genomes of cyprinoid fish including 30 freshwater and two extremophile fish species, *L. waleckii* and *G. przwalskii*, representing two independent origins of extremophiles. We utilized comparative genomics to test whether convergent adaptation to extreme environment in both extremophile lineages is associated with 1) convergent genome-wide patterns of molecular evolution; 2) convergent shifts in selections for specific genes; and 3) convergence in gene expression changes.

## Methods and Materials

### Data processing

We collected the whole genomes of cyprinoid fishes from online databases, NCBI (https://www.ncbi.nlm.nih.gov/genome) and CNGBdb (https://db.cngb.org/) (updated by January 2022) (Table S1). We then applied the BUSCO pipeline (Simão *et al*. 2015) to assess the completeness of cyprinoid fish genome contents based on the Actinopterygii_odb10 single-copy orthologous gene set from OrthoDB (https://www.orthodb.org) that included 3,640 conserved genes. We executed strict criteria to retain the genomes with both genome assemblies and annotated protein-coding genes covering more than 90% conserved genes for further analyses.

### Genome reassembly and reannotation

The genome of *L. waleckii* (GCA_900092035.1) was previously sequenced and assembled (Xu *et al*. 2017) but with relatively low quality (BUSCO: 74.0% completed Actinopterygii conserved genes) compared with the other high-quality cyprinoid fish genomes (BUSCO: > 90% completed Actinopterygii conserved genes) involved in our study. We sought to improve the quality of genome assembly and reassemble the genome of *L. waleckii*. Specifically, we trimmed the adaptors and low-quality sequencing reads using Trimmomatic (Bolger *et al*. 2014). We then *de novo* assemble a draft genome using SPAdes (Nurk *et al*. 2013) and improved scaffolds using RagTag (https://github.com/malonge/RagTag) with a reference chromosome-level genome of *Leuciscus idus* (GCA_021554675.1). We attempted to further improve the scaffolds using Redundans (https://github.com/Gabaldonlab/redundans) with recently released sequencing data (Wang *et al*. 2021), L_RNA_scaffolder (https://github.com/CAFS-bioinformatics/L_RNA_scaffolder) with transcriptome sequencing data (Xu *et al*. 2013a; b), and PEP_scaffolder (https://github.com/CAFS-bioinformatics/PEP_scaffolder) with annotated cyprinoid fish protein datasets (Table S1). Finally, we used BUSCO to assess the completeness of genome reassembly as previously described.

We took advantage of the combination of homology-based and *de novo* prediction approaches to annotate the newly reassembled *L. waleckii* draft genome and other draft cyprinoid fish genomes (Table S1). Specifically, we used AUGUSTUS (Stanke *et al*. 2006) with zebrafish (*Danio rerio*) as the training set for *de novo* annotation. For homolog-based prediction, we download protein sequences of annotated cyprinoid fish genomes from NCBI (Table S1). The candidate genes were firstly identified by aligning these protein sequences to draft genomes using BLAT (github.com/djhshih/blat). We used the MAKER pipeline (Cantarel *et al*. 2008) to annotate the draft genomes (Fig. 1). Finally, we integrated the gene models predicted by both approaches using GLEAN (Elsik *et al*. 2007) with default parameters to remove redundant genes.

### Ortholog identification

We used the OMA (Altenhoff *et al*. 2021) pipeline to infer orthologous groups (OGs) among 32 cyprinoid fish genomes. We included the protein-coding gene dataset from each fish genome and pooled them into a local protein database. We then conducted paralleled all-against-all alignments using DIAMOND (https://github.com/bbuchfink/diamond). We identified the putative OGs from the alignments using OMA with the default setting. Moreover, we further identified one-to-one, one-to-many, and many-to-many orthologs among these fish species. For each 1:1 orthologous pair, we selected the longest transcripts associated with recorded OGs for each species as putative single-copy orthologs. Gene ontology (GO) terms were assigned to single-copy orthologs using Trinotate (https://github.com/Trinotate).

### Genome-scale phylogeny construction and divergence time estimation

We used a phylogenomic approach to reconstruct the phylogeny based on large-scale single-copy orthologs. Specifically, we prepared a dataset that included the amino acid (AA) sequences of 3,784 single-copy orthologs shared by 32 cyprinid fish species and one outgroup species, *Oryzias latipes*. Then, we performed AA alignments using MUSCLE (https://github.com/rcedgar/muscle) and trimmed gaps using trimAL (https://github.com/inab/trimal). In addition, we executed strict criteria to retain the AA alignments of single-copy orthologs with a minimum of 200 AA in length. We assembled a concatenated dataset including all core shared single-copy orthologs and detected the best-fit model of sequence evolution using ModelFinder (Kalyaanamoorthy *et al*. 2017) and reconstructed the maximum likelihood (ML) phylogenetic tree using RAxML (Stamatakis 2014). Finally, we performed the divergence time estimation using MCMCtree in PAML package (Yang 2007) with the topology of 4DTV position and recorded calibration time based on the Timetree (http://www.timetree.org/).

### Molecular convergence analysis

We prepared the codon alignments of 17,410 single-copy orthologs, derived from AA alignments and the corresponding DNA sequences using PAL2NAL v.14 (www.bork.embl.de/pal2nal). We estimated two discrete categories of non-synonymous to synonymous rate ratio (i.e., dN/dS) for extremophile and non-extremophile fish taxa with the concatenated codon sequences of 3,784 core shared orthologs (genome-wide) using HyPhy 2.5 (Kosakovsky Pond *et al*. 2020a) (Fig. 1). We then employed a likelihood ratio test for the comparison of genome-wide dN/dS. To distinguish the evolutionary driving force during the convergent phenotypic transition from non-extremophile to extremophile, we used RELAX (Wertheim *et al*. 2015) to detect the signal of relaxed or intensified selection at genome-wide scale with the concatenated codon sequence datasets and applied a likelihood ratio test.

We sought to determine whether specific genes experience convergent shifts in selective pressures, including convergent acceleration and positive selection in extremophile relative to non-extremophile fish taxa. For convergent acceleration, we estimated the relative evolutionary rate (RER) using the R package, RERconverge (Kowalczyk *et al*. 2019), which calculated relative branch lengths by normalizing branches for focal branches (i.e., extremophile fish branches) to the distribution of branch lengths across all genes, as previously described (Kowalczyk *et al*. 2020; Xu *et al*. 2021) (Fig. 1). Specifically, we prepared AA alignments of 17,410 single-copy orthologs shared by up to 33 fish species. We estimated the branch lengths of each gene tree based on the inferred species tree using the R package, phangorn (Schliep 2011), and calculated RER for each branch within the corresponding gene tree. To correlate RERs with phenotypes, we applied a correlation analysis over the binary variable of focal foreground branches (i.e., extremophile fish branches) and employed a Wilcoxon rank-sum test. The genes with significantly higher RERs (*p* < 0.05) in extremophile fish taxa and positive Rho values as rapidly evolving genes. In addition, we applied gene set enrichment analysis for the gene list generated from the correlation analysis using the fastwilcoxGMT and fastwilcoxGMTall functions in RERconverge (Kowalczyk *et al*. 2019).

For convergent positive selection, we integrated three approaches based on the branch-site (BS) model test to identify genes under positive selection (i.e., positively selected gene, PSG) in extremophile fish taxa, including aBS-REL (Smith *et al*. 2015), BUSTED (Murrell *et al*. 2015), and Contrast-FEL (Kosakovsky Pond *et al*. 2020b) (Fig. 1). In brief, we prepared codon alignments of 17,410 single-copy orthologs shared by up to 33 fish species. We performed the BS models tests for each gene with codon alignment and pruned gene tree, and then applied the corresponding statistical adjustments as previously described (Murrell *et al*. 2015; Smith *et al*. 2015; Kosakovsky Pond *et al*. 2020b). Finally, we performed the gene ontology enrichment analysis for the PSG repertoire using the R package, topGO (https://rdrr.io/bioc/topGO).

### Protein modelling

We sought to detect convergence in amino acid sites on a specific gene that experienced positive selection, that is convergent positively selected site (PSS). To determine the potential functional effect of convergent PSS in a specific gene, we applied a protein structure homology-modelling approach to generate three-dimensional (3D) protein structure of candidate gene using a protein structure prediction tool, I-TASSER (https://zhanggroup.org/I-TASSER/). We finally mapped PSS onto the predicted 3D protein structure using PyMOL 2.4 (https://pymol.org/2/).

### Comparative transcriptomics

We retrieved tissue-specific transcriptome sequencing data (gill, kidney, and liver) for *G. przwalskii* (Li *et al*. 2021) and *L. waleckii* (Xu *et al*. 2013b; Chen *et al*. 2019) in freshwater and extreme envrironments from NCBI SRA database (https://www.ncbi.nlm.nih.gov/sra). Specifically, we collected the tissue-specific transcriptomes for *G. przwalskii* that sampled from Lake Qinghai (extreme envrionment) and Quanji River (freshwater), *L. waleckii* that sampled from Lake Dali Nur (extreme environment) and Shali River (freshwater). We then proceeded the quality control and removed low-quality reads using Trimmomatic as previously described. We further mapped all reads to protein-coding genesets of two extremophile fish species using kallisto (https://pachterlab.github.io/kallisto) to obtain expected counts and transcripts per million (TPM) for each gene. Finally, we performed differential gene expression analysis by comparing three tissues of fish individuals from extreme saline/alkaline and freshwater environments using R package, DESeq2 (Love *et al*. 2014) (Fig. 1).

## Results

### Genomes, phylogeny, and independent origins of extremophile fishes

We initially obtained genome data of 43 fish species from 5 major families within suborder Cyprinoidei, including Cyprinidae, Leuciscidae, Xenocyprididae, Danionidae, Gobionidae (Table S1). In addition, we reassembled and improved the genome of *L. waleckii* with recently sequenced *L. waleckii* transcriptomes, chromosome-level genome of closely related species (*L. idus*), and annotated cyprinoid protein datasets (Table S1). We assessed the completeness of genome contents of these fish species using BUSCO (https://busco.ezlab.org) and retained a total of 32 high-quality genomes covering more than 90% Actinopterygii conserved genes (3,640 genes). We further used a combination of homology and *de novo* approaches to annotate the gene models of reassembled *L. waleckii* genome and other unannotated cyprinoid genomes, resulting in gene models ranged from 23,906 to 96,703 per species (Table S1) with an average of 95.85% of BUSCO Actinopterygii conserved genes.

We used OMA pipeline (Altenhoff *et al*. 2021) to obtain 17,410 orthologous groups (OGs) and 3,784 core single-copy orthologs (one to one) that included all 32 cyprinoid species (Table S2). We reconstructed a genome-scale phylogeny based on the concatenated core single-copy ortholog data, which was strongly supported for all nodes with 100% bootstrap value, which pinpointed two independent origins of extremophile in cyprinoids (Fig. 1A). We further estimated divergence time for each phylogenetic node. The divergence time between two *Gymnocypris species* was estimated to be less than one million years ago, which suggested a recent origin of extremophiles in cyprinoids (Fig. 1A).

### Convergence in genome-wide patterns of molecular evolution in extremophile fish taxa

First, we ask whether extremophile fish taxa had convergent genome-wide patterns of molecular evolution (Non-synonymous/Synonymous rate ratio, dN/dS) compared to non-extremophile fish taxa. We further took advantage of the HyPHY pipeline (Kosakovsky Pond *et al*. 2020a) to test the hypotheses by comparing dN/dS between *a priori* defined extremophile fish branches (focal foreground branches) and non-extremophile fish branches (background branches) across the phylogeny with concatenated single-copy ortholog data. We found that extremophile fish taxa exhibited significantly higher dN/dS than non-extremophile fish taxa (Likelihood ratio test, *p* < 0.00001) (Fig. 1B). Elevation in dN/dS can be caused by intensified positive selection or relaxed purifying selection. To determine which evolutionary driving force acting during the convergent phenotypic transition to extremophile, we detected the signal of intensified selection in extremophile relative to non-extremophile fish taxa (Likelihood ratio test, *p* < 0.00001, k = 1.07) using RELAX (Wertheim *et al*. 2015). In detail, RELAX models the distribution of three categories of dN/dS — positive selection (dN/dS > 1), neutral evolution (dN/dS = 1), purifying selection (dN/dS < 1) — across a specified phylogeny, comparing focal foreground (i.e., extremophile) to background (i.e., non-extremophile) branches and estimating K value that indicates overall relaxation (k < 1) or intensification (k > 1). Our results further showed that the convergent phenotypic transition to extremophile is caused by an overall intensification of selection across the whole genome, which is strengthening both purifying selection and positive selection away from neutrality (Fig. 1C, Table S3).

### Convergent acceleration in gene-wide rates of molecular evolution in extremophile fish taxa

Next, we ask if there was evidence for convergence in gene-wide rates of molecular evolution, as we observed convergence in genome-wide acceleration in extremophile fish taxa. We utilized RERconverge (Kowalczyk *et al*. 2019), an R package specifically developed to test for the signature of convergent molecular evolution underlying convergent phenotypic evolution. RERconverge examines the rate of protein sequence evolution (i.e., relative evolutionary rate, RER) for each gene on every branch across the specified phylogeny, standardized to the distribution of rates across all genes. Genes for which these standardized rates are consistently either higher or lower in focal foreground branches (i.e., extremophile fish branches) compared to background branches (i.e., all remaining branches) are identified as having experienced acceleration in rate of molecular evolution in extremophile fish branches. By analyzing all 17,410 shared orthologs, we identified 112 genes experiencing convergent acceleration in extremophile fish branches (Wilcoxon rank sum test, *p* < 0.05), including calcium channel voltage-dependent gamma 7 subunit (CACNG7), immune-related lectin-like receptor 1 (ILLR1), potassium voltage-gated channel subfamily S member 2 (KCNS2), sodium- and chloride-dependent taurine transporter (SLC6A6), solute carrier family 2, facilitated glucose transporter member 1 (SLC2A1) (Fig. 3A, Table S4). We also identified 27 enriched gene ontology (GO, Biological Processes) terms under convergent acceleration in extremophile fish branches (Benjamini-Hochberg adjusted *p* < 0.05) (Fig. 3B, Table S4). These significantly enriched GO terms were mainly related to nervous system development including synaptic transmission (GO:0007268), neurotransmitter secretion (GO:0007269), memory (GO:0007613), ion transport including calcium ion import (GO:0070509), potassium ion transmembrane transport (GO:0071805), hormone response including cellular response to hormone stimulus (GO:0032870), and immune response including homophilic cell adhesion (GO:0007156) (Fig. 3B, Table S5).

**Figure 3.**
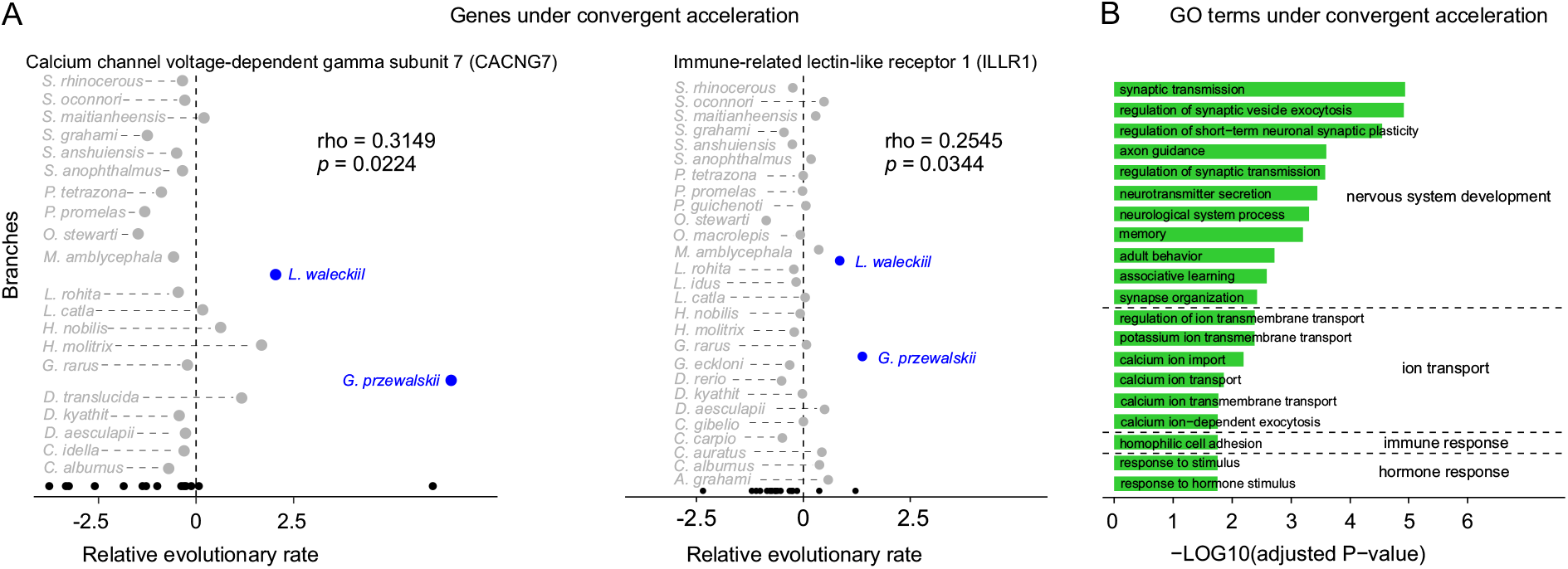
Genes and GO terms under convergent acceleration in both extremophile fishes. (A) Two representative cases of genes under convergent acceleration in extremophile fish branches relative to non-extremophile fish branches. One is calcium channel, voltage-dependent, gamma subunit 7a (CACNG7), another is immune-related lectin-like receptor 1 (ILLR1). Extremophile fish branch, non-extremophile fish branch, and ancestral branch were represented by grey, blue, and dark dots, respectively. (B) Bar plot depicting the significantly enriched GO terms under convergent acceleration (Benjamini-Hochberg adjusted *p* < 0.05), including four major functional categories, nervous system development, ion transport, immune response, and hormone response.

### Convergent signatures of positive selection at the gene level in extremophile fish taxa

We further ask whether there is evidence for convergent positive selection acting on specific genes during convergent phenotypic transition, as we found the signal of intensified positive selection in extremophile fish taxa. We utilized three complementary approaches based on branch-site model to identify putative genes under positive selection in extremophile fish taxa. Specifically, aBS-REL (adaptive Branch-Site Random Effects Likelihood) (Smith *et al*. 2015) tests if positive selection (dN/dS > 1) has occurred on a proportion of branches across the phylogeny, BUSTED (Branch-Site Unrestricted Statistical Test for Episodic Diversification) (Murrell *et al*. 2015) detects positive selection acting on at least one site on at least one branch, while both models cannot detect positive selection acting on specific sites, alignment-wide. To further test for positive selection at specific individual amino acid site, we used Contrast-FEL (Kosakovsky Pond *et al*. 2020b) to determine whether a specific site in a gene under different selective pressures (e.g. positive selection, dN/dS > 1) in *a priori* defined focal foreground branches (i.e., extremophile fish branches) relative to background branches (i.e., all remaining branches). First, we identified 448 putative positively selected genes (PSGs) in *L. waleckii* and *G. przwalskii* (Holm-Bonferroni corrected *p* < 0.05) by aBS-REL. We then used BUSTED to retain 215 PSGs with at least one site on at least one branch under positive selection (Benjamini-Hochberg adjusted *p* < 0.05). Finally, we used Contrast-FEL to identify 214 core-shared PSGs with specific amino acid sites under positive selection (False discovery rate corrected *p* < 0.05) (Fig. 4A, Table S6). These PSGs were significantly enriched in 184 GO terms (Biological Processes) within four major functional categories (Fisher’s exact test, *p* < 0.05) (Fig. 4B, Table S7), including nervous system development (GO:0051588, regulation of neurotransmitter transport; GO:0099536, synaptic signaling; GO:0007423, regulation of synapse assembly, GO:0007417: central nervous system development, GO:0050807: regulation of synapse organization), and ion transport function (GO:0006821, chloride transport; GO:0071805, potassium ion transmembrane transport; GO:0071470: cellular response to osmotic stress, GO:0070839: metal ion export), immune response (GO:0002922: positive regulation of humoral immune response, GO:0060330: regulation of response to interferon-gamma, GO:0071353: cellular response to interleukin-4), and reproduction (GO:0048599: oocyte development, GO:0009994: oocyte differentiation).

**Figure 4.**
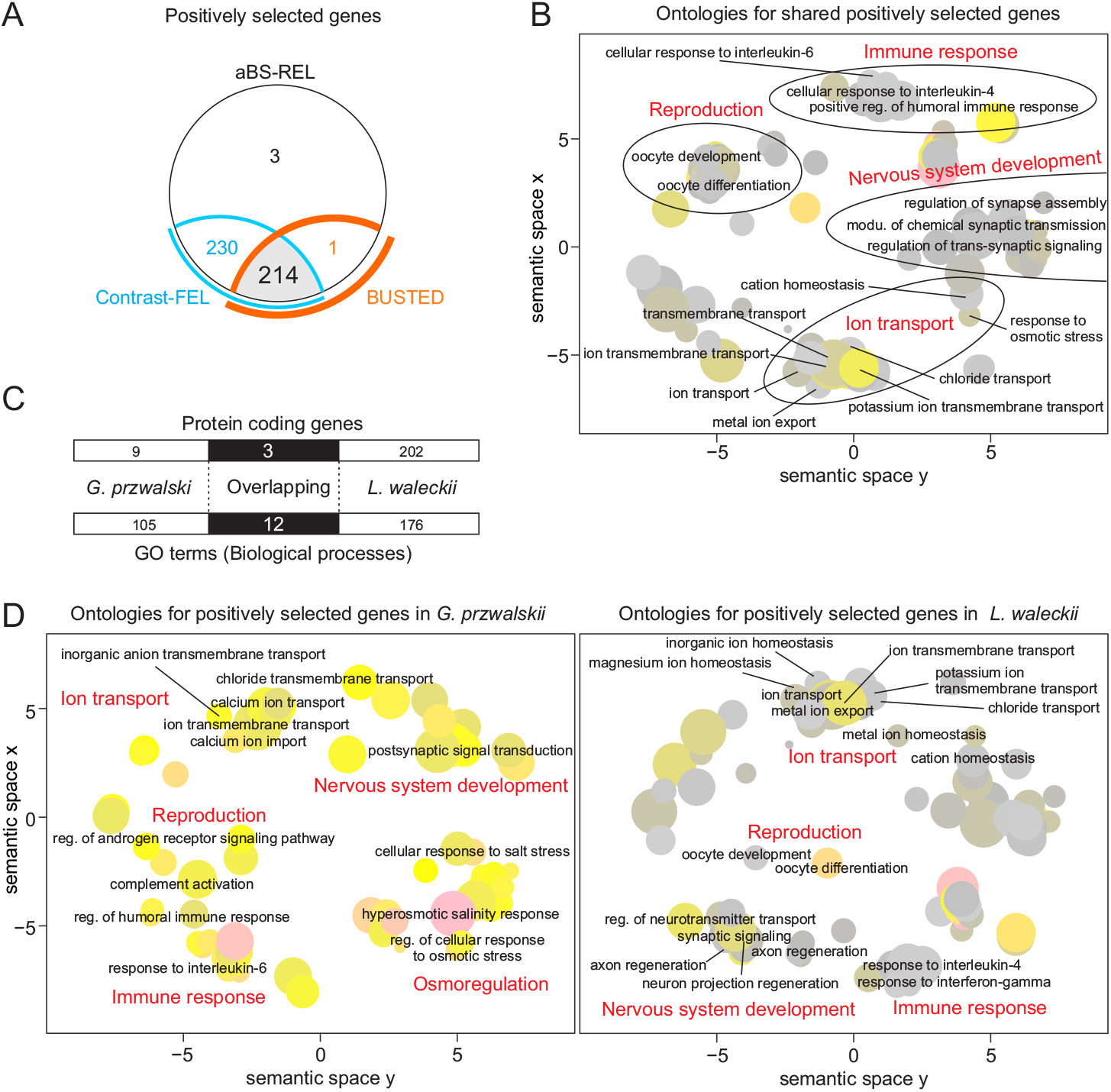
Genes and GO terms under convergent positive selection in extremophile fishes. (A) Venn diagram depicting numbers of PSGs in extremophile fishes identified by three branch-site models, aBSREL, BUSTED and Contrast-FEL. (B) REVIGO plot depicting the dominant enriched GO terms for shared PSGs in both species, including nervous system development, ion transport, reproduction, and immune response. The scale of dot indicating the number of included enriched GO terms under a dominant GO term, the color scale representing the *p* value transformed by log10. (C) The number of unique and shared protein-coding genes and enriched GO terms (Biological process) under positive selection. (D) REVIGO plot depicting the dominant enriched GO terms for PSGs in *G. przwalskii* and *L. waleckii*.

Specifically, most PSGs were lineage-specific and present in either *L. waleckii* or *G. przwalskii*, such as *potassium voltage-gated channel subfamily S member 3* (*KCNS3*), *sodium/calcium exchanger 3* (*NAC3*), *sodium/potassium/calcium exchanger 5* (*NCKX5*), and *solute carrier family 2, facilitated glucose transporter member 11* (*GTR11*) (Fig. 4C, Table S6). Besides lineage-specific PSGs, we found few shared PSGs (N = 3) in both extremophile fish taxa, including *extracellular calcium-sensing receptor* (*CaSR*), *zinc finger MYND domain-containing protein 11* (*ZMYND11*), and *importin subunit alpha-1* (*IMA1*) (Fig. 4C, Table S6). We further applied GO term enrichment analyses for PSGs of each extremophile fish lineage. Similarly, we found both PSG datasets were significantly enriched into four major functional categories (Fisher’s exact test, *p* < 0.05), including nervous system development, ion transport, reproduction, and immune response (Fig. 4D, Table S8), although a small number of overlapping enriched GO terms (N = 12) were present in both extremophile fish taxa, such as regulation of cellular response to stress (GO:0080135) and transmembrane transport (GO:0055085) (Fig. 4D, Table S9).

Additionally, we sought to identify convergent positively selected AA site (PSS) that occurred in both extremophile fish taxa. That is the exact same AA sites in both *L. waleckii* and *G. przwalskii* under positive selection. For this, we inspected AA alignments for all PSGs. We found no convergent PSS in both extremophile fish taxa. Instead, convergent positive selection on the AA site level appears to be relatively “loose”, that is, extremophile fish taxa harbor different PSS on the exact same gene. For example, in a calcium ion transport gene, CaSR (Fig. 5A), we identified a set of “convergent” positively selected AA sites (Fig. 5A). Specifically, we identified 10 unique PSS in *G. przewalskii* and another two unique PSS in *L. waleckiil* (Fig. 5B). We predicted the protein structure of extremophile fish CaSR based on the structure of human calcium-sensing receptor (PDB accession: 7DTT) using homology-modelling approach. Further, we mapped these 12 PSS onto the three-dimensional protein structure of CaSR that located at venus flytrap domain (VFT) (Fig. 5C), which may affect the activity in ion transport by dimerization.

**Figure 5.**
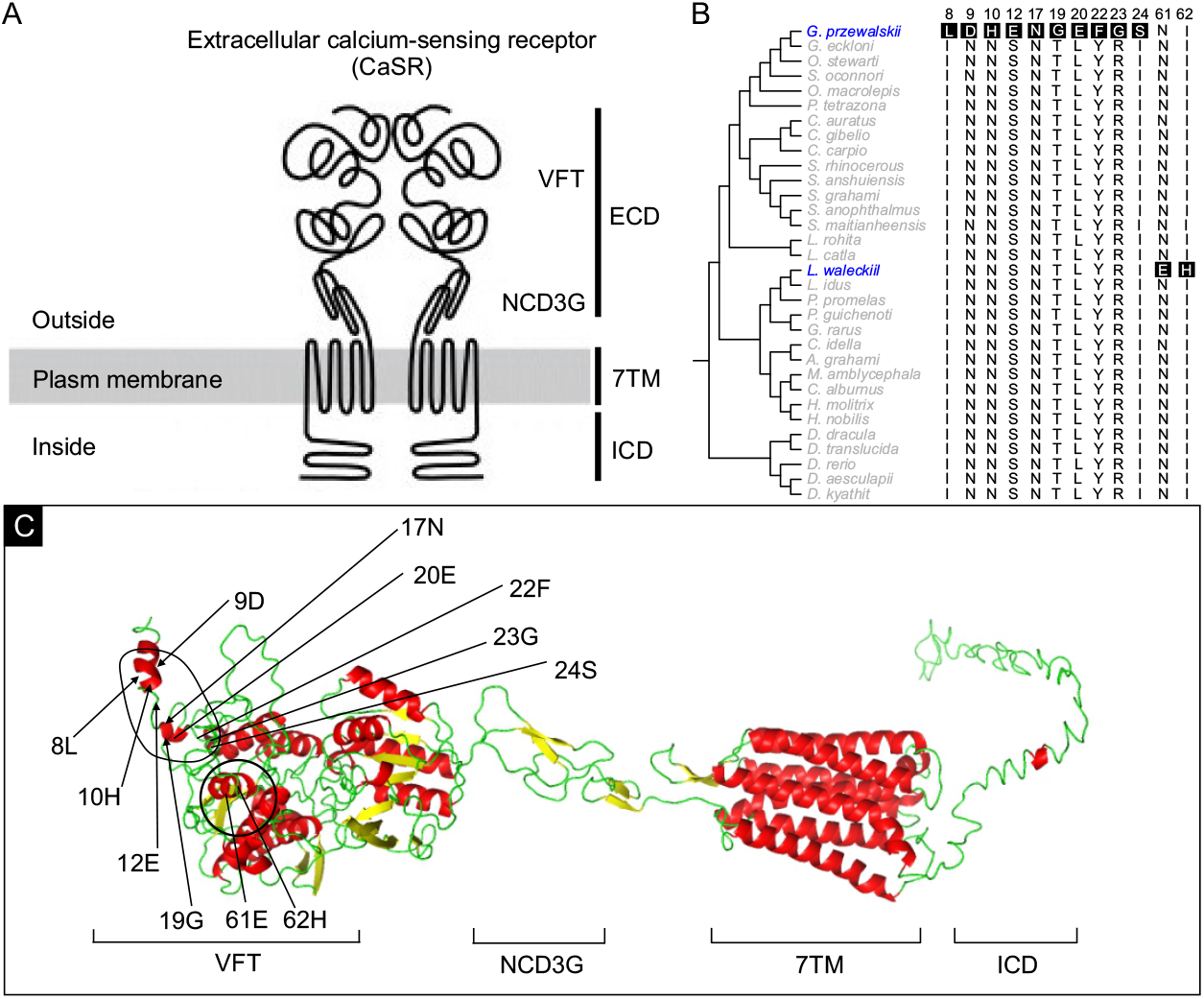
Predicted protein structure for extracellular calcium-sensing receptor (CaSR) in extremophile fish. (**A**) Schematic diagram depicting the topology and domain arrangement in CaSR, including extracellular domain (ECD), seven transmembrane domain (7TM) and intracellular domain (ICD). ECD includes bilobed venus flytrap (VFT) and nine-cysteines domain of family 3 G protein-coupled receptor (NCD3G) modules. Dimerization occurs via both covalent (disulfide) and non-covalent bonds between ECDs of two receptor monomers. (**B**) Positively selected amino acid sites (PSS) in CaSR. Extremophile and non-extremophile fish species name are highlighted by blue and grey, respectively. The PSS in extremophile fish taxa were shadowed by dark boxes. (**C**) Three-dimensional (3D) homology protein structure of *G. przewalskii* CaSR. The PSS residues are labeled, and all located in VFT domain.

### Convergent roles of differentially expressed genes under selection in extremophile fish taxa

Finally, we sought to determine the convergence of changes in gene expression during convergent adaptation to extreme environment in cyprinoid fishes. We compared gill, kidney, and liver tissues-specific transcriptomes between individuals in extremely saline/alkaline and freshwater environments to identify significantly differentially expressed genes (DEGs) (Benjamini and Hochberg, adjusted *p* < 0.05) for *L. waleckii* and *G. przwalskii*, respectively (Fig. 6A). We found a substantial set of common DEGs shared by both extremophile fish taxa in each tissue (Fig. 6B, Table S10). Specifically, we observed 21 common DEGs in gill tissue, such as *ectoderm-neural cortex protein 1* (*ENC1*) and *caspase recruitment domain-containing protein 14* (*CARD14*) involved in immune response, showing consistently up-regulated or down-regulated in extreme environment; a total of 33 DEGs showed similar expression pattern between two species in kidney, such as *aquaporin 1* (*AQP1*) and *solute carrier family 41 member 1* (*SLC41A1*) involved water and ion transport process; we also found 17 common DEGs in liver tissue, such as *apolipoprotein C-II precursor* (*APOC2*) associated with lipid metabolism and *fibrinogen gamma chain precursor* (FGG) associated with immune response (Table 1). To further identify genes that were differentially expressed and experienced convergent shift in selections, we examined the overlapping results from differentially expressed gene sets and rapidly evolving / positively selected gene sets. We found a few rapidly evolving genes (N = 5) were significantly differentially expressed in *G. przwalskii*, such as *sodium- and chloride-dependent taurine transporter* (*SLC6A6*) associate with ion transport function, *OTU domain-containing protein 5* (*OTUD5*) associated with innate immune response (Table 1). Similarly, we found a small number of genes (N = 11) with evidence for both selection and differential expression in *L. waleckii*, such as *heat shock protein HSP 90-alpha 1* (*HSP90AA1*) associated with reproduction, *transcription factor CP2-like protein 1* (*TFCP2L1*) that involved in acid-base and salt-water homeostasis. In addition, we observed a few rapidly evolving genes were differentially expressed in both kidney and liver tissues, such as an ion transport gene, *polycystic kidney disease protein 1-like 2* (*PKD1L2*), and another gene, *DNA helicase MCM8* (*MCM8*) associated with reproduction (Table 1).

**Figure 6.**
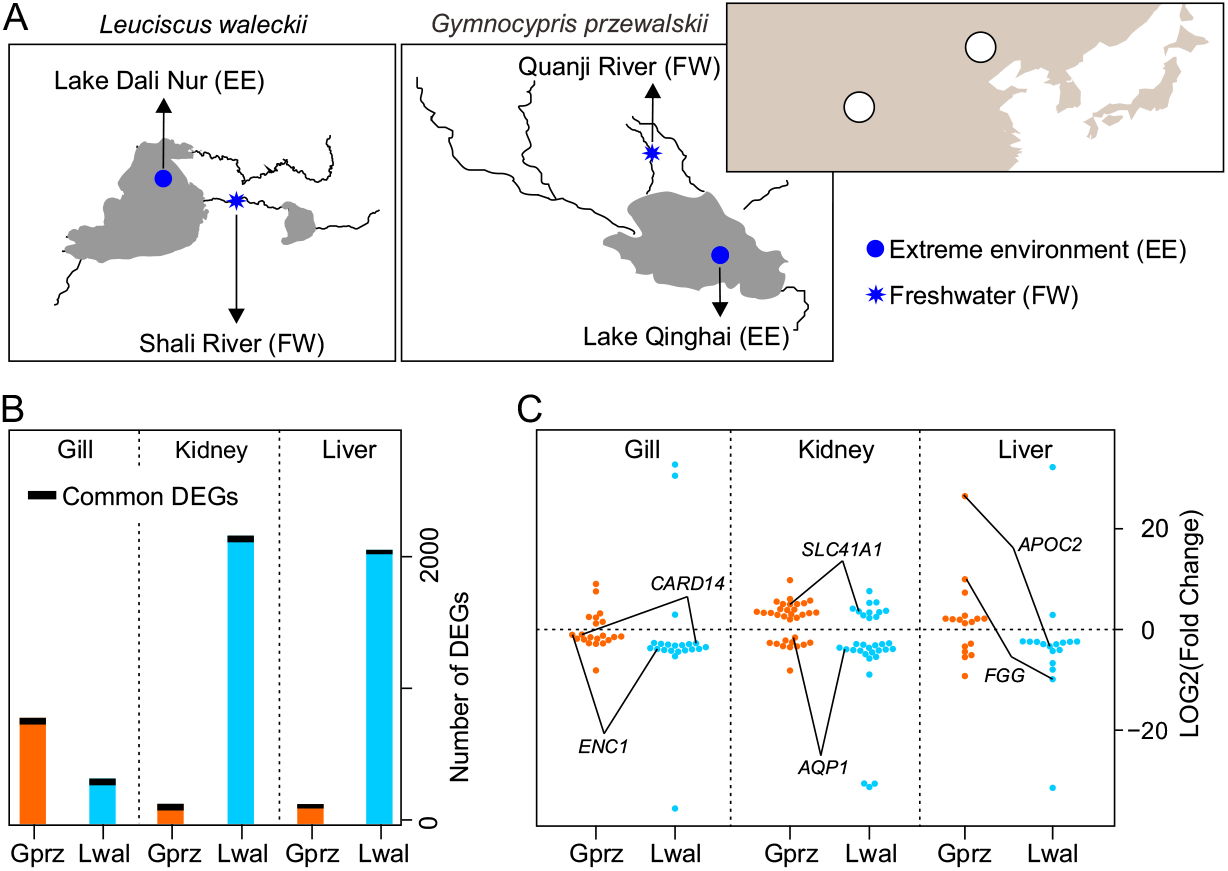
Convergence in changes of gene expressions in extremophile fish taxa. (A) Map showing the geographic locations of *Leuciscus waleckii* from the Lake Dali Nur and the Shali River, *Gymnocypris przewalskii* from the Lake Qinghai and the Quanji River. Blue dot representing the extremely saline and alkaline environment, and blue star dot represents the freshwater environment. (B) Bar plot depicting the number of differentially expressed genes (DEGs) between freshwater and extremely saline / alkaline environment for each species. Orange and light blue bars represent the DEGs in gill, kidney, and liver tissues of each species. Dark bar represents the overlapping DEGs (also known as, common DEGs) between extremophile fish taxa in each tissue. (C) Dot plot depicting the expression patterns of common DEGs between two extremophile fish taxa, including upregulation (LOG2Fold-change > 0) or downregulation (LOG2Fold-change < 0).

**Table 1.** Representative differentially expressed genes under convergent selections in extremophile fish taxa.

| DEG name | Gene description | Species | Expressed tissue | Functional category |
| --- | --- | --- | --- | --- |
| Genes under convergent acceleration |  |  |  |  |
| *SLC6A6* | Sodium- and chloride-dependent taurine transporter | *G. przewalskii* | Gill | Ion transport |
| *A2ML1* | Alpha-2-macroglobulin-like protein 1 | *G. przewalskii* | Kidney | Immune response |
| *OTUD5* | OTU domain-containing protein 5 | *G. przewalskii* | Liver | Immune response |
| *HSP90AA1* | Heat shock protein HSP 90-alpha 1 | *L. waleckii* | Gill | Reproduction |
| *APC2* | Adenomatous polyposis coli protein 2 | *L. waleckii* | Gill | Nervous system development |
| *MPEG1* | Macrophage expressed 1 | *L. waleckii* | Kidney | Immune response |
| *PKD1L2* | Polycystic kidney disease protein 1-like 2 | *L. waleckii* | Kidney, Liver | Ion transport |
| *MCM8* | DNA helicase MCM8 | *L. waleckii* | Kidney, Liver | Reproduction |
| *MTMR7* | Myotubularin-related protein 7 | *L. waleckii* | Kidney, Liver | Reproduction |
| *TFCP2L1* | Transcription factor CP2-like protein 1 | *L. waleckii* | Liver | Acid-base balance |
| Genes under positive selection |  |  |  |  |
| *RTN4RL2* | Reticulon-4 receptor-like 2 precursor | *G. przewalskii* | Gill | Brain development |
| *CRB2* | protein crumbs homolog 2 | *G. przewalskii* | Kidney | Nervous system development |
| *PDE6C* | Cone cGMP-specific 3',5'-cyclic phosphodiesterase subunit alpha' | *G. przewalskii* | Liver | Ion transport |
| *NOTCH1* | Neurogenic locus notch homolog protein 1 | *L. waleckii* | Gill | Nervous system development |
| *NCAM1* | Neural cell adhesion molecule 1 | *L. waleckii* | Gill, Kidney, Liver | Nervous system development |
| *OSBPL10* | Oxysterol-binding protein-related protein 10 | *L. waleckii* | Kidney | Lipid metabolism |
| *EFHD1* | EF-hand domain-containing protein D1 | *L. waleckii* | Kidney, Liver | Ion transport |
| *PCDH18* | Protocadherin-18 | *L. waleckii* | Kidney, Liver | Ion transport |
| *KPNA2* | Importin subunit alpha-1 | *L. waleckii* | Liver | Energy metabolism |
| *PAMR1* | Inactive serine protease PAMR1 | *L. waleckii* | Liver | Immune response |

Besides rapidly evolving DEGs, we aslo identified a set of DEGs under positive selection (Table 1). We found 5 positively selected DEGs in *G. przwalskii*, such as *reticulon-4 receptor-like 2 precursor* (*RTN4R2*) and *protein crumbs homolog 2* (*CRB2*) that involved in nervous system development. Moreover, we found 15 positively selected DEGs in *L. waleckii*, such as *neurogenic locus notch homolog protein 1* (NOTCH1) and, *EF-hand domain-containing protein D1* (*EFHD1*) which associated with nervous system development, an ion transport gene, *protocadherin-18* (*PCDH18*), and *inactive serine protease PAMR1* (*PAMR1*) that involved in immune response (Table 1).

## Discussion

Our results support the three hypotheses and identify common genomic signatures associated with convergent adaptation to extreme environment in cyprinoid fish. Specifically, we found: 1) convergently accelerated genome-wide rate of nonsynonymous substitution and signal of intensified positive selection in extremophile fish lineages; 2) hundreds of genes tended to experience convergent acceleration in rates of nonsynonymous substitution and convergent positive selection in extremophile fish lineages; 3) a substantial set of common differentially expressed genes under convergent selection in gill, kidney and liver tissues of extremophile fish. These genes were associated with several key functions, such as nervous system development, reproduction, ion transport and immune response. Altogether, these findings suggest that extremophile fishes may convergently evolve to adapt to extreme environments in a concerted way through both sequence changes and expression shifts.

### Enhanced nervous systems

Extreme environments have impacted the physiology of fish in a wide variety of ways (Evans & Claiborne 2005). Although recent advances in fish genomics have revealed multiple genetic adaptations in extremophile fishes (Wang & Guo 2019), very few studies have indicated the association between extreme-environment adaptation and the evolution of genes that involved in nervous system development. A recent comparative transcriptomic study had identified a set of microtubule regulation related genes under positive selection that regulate axon formation, and suggested deep-sea fish may have more developed nervous systems to adapt to high hydrostatic pressure environments (Lan *et al*. 2018). Intriguingly, our genome-wide analysis identified a set of nervous system development associated genes that experienced convergent acceleration or convergent positive selection in two extremophile fish lineages. For example, the gene *calcium channel voltage-dependent gamma 7 subunit* (*CACNG7*), which experienced acceleration in *G. przwalskii* and *L. waleckii*, regulates synaptic transmission, and mutation in CACNG7 has been associated with multiple neurological disease (Burgess & Noebels 1999). The gene *reticulon-4 receptor-like 2* (*RTN4RL2*), that showed differentially expressed in response to extremely saline and alkaline in *G. przwalskii* and also experienced positive selection, functions in axon migration and neuron projection development (Wills *et al*. 2012). Although we found a small number of restrictedly overlapping differentially expressed genes under convergent selections in both extremophile fish lineages, different genes involved in similar conserved nervous system development pathways were confirmed, indicating that these genes may share similar evolutionary paths in convergent phenotypic transition to extremophile. Collectively, this finding provides putative genomic evidence for enhanced nervous systems contributing to extremophile fish in convergent adaptation to extreme environments, such as extremely saline and alkaline water.

### Rapidly evolving reproduction system

Extreme environments have disruptive effects on endocrine system of fish, can interfere their reproductive functions (Wang & Guo 2019). Notably, extremophile fish *G. przwalskii* and *L. waleckii* are both anadromous and migrate to spawn in rivers (freshwater) (Xu *et al*. 2017; Tong *et al*. 2017, 2021). Reproduction in most anadromous fishes is a seasonal phenomenon to ensure the survival of their offspring, which fish experienced the transition from saline or alkaline water to freshwater (Tian *et al*. 2019). This spawning migration behavior may require their reproduction system rapidly adapted to the environmental transition. Indeed, our study identified a set of common differentially expressed genes under convergent acceleration in both extremophile fish, as well anadromous fish. For example, the gene *heat shock protein HSP 90-alpha 1* (*HSP90AA1*), regulates spermatogenesis and has the association with spawning migration in another anadromous fish, *Coilia nasus* (Fang *et al*. 2016). Another differentially expressed gene *DNA helicase MCM8* (*MCM8*), also experienced convergent acceleration, is essential for gametogenesis, and mutation in MCM8 has been associated with ovarian failure (AlAsiri *et al*. 2015). In addition, we also found key GO terms related to reproduction were under convergent positive selection in extremophile fish. It is possible that rapid evolution of reproduction related genes may facilitate convergent adaptation to extreme environments in anadromous fish.

### Convergent evolution of ion transport, osmoregulation, and acid-base balance functions

Extreme environments, such as highly salinity and alkalinity are challenging the osmoregulation system of fish. During the transition from freshwater to highly saline or alkaline water, fish must increase its rate to balance osmotic loss of water to more solute-concentrated environment and actively excrete Na^+^ and Cl^−^ from gill to maintain ionic and osmoregulation (Marshall 2005). Thus, extremophile fish require enhanced physiological abilities including ion transport, osmoregulation, and acid-base balance to respond the increase in salinity and alkalinity of freshwater (Marshall 2005; Evans *et al*. 2005). Past studies, including ours, have identified key genes associated with osmoregulation and ion transport in *G. przwalskii* and *L. waleckii* under rapid evolving (aka., acceleration) and positive selection (Xu *et al*. 2017; Tong *et al*. 2017, 2021; Tong & Li 2020). Consistently, we identified a substantial set of ion transport, osmoregulation, and acid-base balance related genes are under convergent selection in both extremophile fish lineages. For example, the genes in solute carriers family (SLC), that encoded transmembrane transporters for inorganic ions, amino acids, neurotransmitters, sugars, purines and fatty acids, and other solute substrates (Dorwart *et al*. 2008), such as *solute carrier family 41 member 1* (*SLC41A1*), *sodium- and chloride-dependent taurine transporter* (*SLC6A6*). In addition, we identified numbers of common differentially expressed genes in tissues of both fish lineages. For example, the gene *aquaporin 1* (*AQP1*), which has been studied extensively in fish gills adaptation to highly saline water (Riesch *et al*. 2015). Besides the molecular convergence at gene-wide level, we also attempted to identify convergence at amino acid site level and suggest a “loose” pattern of site-specific convergence. One of representative cases, the gene *extracellular calcium-sensing receptor* (*CaSR*), regulates the calcium ion homeostasis (Loretz 2008). We observed different positively selected sites located on the same candidate gene, which may cooperatively lead the phenotypic convergence in fish. In the combination of our present results and previous findings, the gene-wide and loose site-wide convergent of genes involved in ion transport, osmoregulation, and acid-base balance are essential to fish convergent adaptation to extreme environment.

### Adaptive Immunity system

Extremely saline and alkaline stress may cause extensive damage to fish immune system (Bowden 2008; Sridhar *et al*. 2020). In *G. przwalskii*, our previous genome-wide studies had identified a set of immune-related genes that experienced rapidly evolving or positive selection (Tong *et al*. 2017, 2021; Tong & Li 2019), while there is no signal of selections on immune-related genes in *L. waleckii* (Xu *et al*. 2017). Although previous studies seem to suggest inconsistent patterns of molecular evolution on immune-related genes in extremophile fish adaptation to extreme environment, our present study provide the evidence of molecular convergence on immune-related genes in both fish lineages relative to other freshwater cyprinoid fishes. For example, the gene *immune-related lectin-like receptor 1* (*ILLR1*), which experienced convergent acceleration, plays roles in fish innate immunity (Yang 2009). Another gene *fibrinogen gamma chain precursor* (*FGG*), which is one of the common differentially expressed genes, facilitates the antibacterial immune response in aquatic organisms (Gordy *et al*. 2015). Besides, we also identified a set of GO terms under convergent acceleration or convergent positive selection in both extremophile fish lineages. Thus, this finding indicates that the convergence in genes sequence changes and gene expression shifts may contribute to fish convergent adaptation to extremely saline and alkaline environment. Overall, our comparative genomic analyses of cyprinoid fish genomes identify common changes in gene sequences and expressions putatively associated with convergent adaptation to extreme environment. Further study should include large scale of genomic data by including more extremophile fish lineages on multiple regulatory levels to unveil the genetic basis of convergent adaptation to extreme environments.

## Data availability

All the scripts required to perform the analyses are available at https://github.com/jiyideanjiao/convergent_adaptation_cyprinoid_fishes.

## Acknowledgements

This work was supported by the Field Research Fund of the University of Pennsylvania Biology Department to C.T.

## Author contributions

C.T. conceived and designed this project. C.T. and M.L. performed the comparative genomics analyses. C.T. wrote the paper. All authors read and approved the final manuscript.

